# Modelling cancer progression using Mutual Hazard Networks

**DOI:** 10.1101/450841

**Authors:** Rudolf Schill, Stefan Solbrig, Tilo Wettig, Rainer Spang

## Abstract

**Motivation:** Cancer progresses by accumulating genomic events, such as mutations and copy number alterations, whose chronological order is key to understanding the disease but difficult to observe. Instead, cancer progression models use co-occurence patterns in cross-sectional data to infer epistatic interactions between events and thereby uncover their most likely order of occurence. State-of-the-art progression models, however, are limited by mathematical tractability and only allow events to interact in directed acyclic graphs, to promote but not inhibit subsequent events, or to be mutually exclusive in distinct groups that cannot overlap.

**Results:** Here we propose Mutual Hazard Networks (MHN), a new Machine Learning algorithm to infer cyclic progression models from cross-sectional data. MHN model events by their spontaneous rate of fixation and by multiplicative effects they exert on the rates of successive events. MHN compared favourably to acyclic models in cross-validated model fit on four datasets tested. In application to the glioblastoma dataset from The Cancer Genome Atlas, MHN proposed a novel interaction in line with consecutive biopsies: *IDH1* mutations are early events that promote subsequent fixation of *TP53* mutations.

**Availability:** Implementation and data are available at https://github.com/RudiSchill/MHN.

## 1 Introduction

Tumours turn malignant in an evolutionary process by accumulating genetic mutations, copy number alterations, and changes in DNA methylation. Such progression events arise randomly in tumour cells, but due to unknown epistatic interactions they tend to fixate in specific chronological orders. Whether an event in- or decreases the reproductive fitness of a tumour cell relative to competing clones depends on preceding events in this cell: a new mutation in a driver gene can be advantageous for the cell in one genomic background and it can be neutral or lethal in another, thus setting the course for the tumour’s future genomic progression.

While progression is a dynamic process, available genotype data are cross-sectional and combine static snapshots from different tumours at different stages of development. Nevertheless, assuming that the tumour genomes are observations from the same stochastic process, cancer progression models can infer epistatic interactions between events from their co-occurence patterns.

To this end, increasingly complex models and learning algorithms have been developed. Fearon and Vogelstein (1990) manually inferred that colorectal cancers progress along a linear chain of mutations in the genes *APC → K-RAS → TP53*. Desper et al. (1999) formalized and extended this concept to Oncogenetic Trees, where a single event can promote multiple successor events in parallel. Beerenwinkel et al. (2007) further generalized these to Conjunctive Bayesian Networks (CBN), where events may require multiple precursors to convey a selective advantage and interactions are hence described by a directed acyclic graph. Other models include Bayesian Networks with different types of acyclic interactions (Farahani and Lagergren, 2013; Misra et al., 2014; Ramazzotti et al., 2015) and networks with cycles (Hjelm et al., 2006) where events can be mutually promoting but not exclusive.

Mutual exclusivity of events, however, is a frequently observed phenomenon in cancer (Yeang et al., 2008). Two events are considered mutually exclusive if they co-occur less frequently than expected by chance. There are at least two mechanisms that can cause this data pattern: (a) the events are synthetically lethal and (b) the events disrupt the same molecular pathway such that whichever event occurs first conveys most of the selective advantage and decreases selective pressure for the others.

Mutual exclusivity is a cyclic interaction between events and thus cannot be naturally encoded by an acyclic model. The currently prevalent workaround was introduced by Gerstung et al. (2011) who first grouped events into pathways and in a second step learned acyclic models on the coarser resolution of pathways. Pathways can either be derived from biological knowledge, learned from data by testing groups of events for mutual exclusivity (Leiserson et al., 2013; Szczurek and Beerenwinkel, 2014; Constantinescu et al., 2015), or by a combination of both (Ciriello et al., 2011; Kim et al., 2015). Raphael and Vandin (2015) pointed out that inferring pathways separately from their interactions can lead to inconsistencies in the presence of noise and presented the first algorithm that simultaneously groups events into pathways and arranges the pathways in a linear chain. PathTiMEx (Cristea et al., 2017) generalizes this from linear chains to acyclic progression networks (CBN).

This approach, however, relies on the strong and unproven assumption that the future evolution of a tumour does not depend on which specific event in a group of mutually exclusive events actually occurred. In fact, we will show below that this interchangeability assumption is not always in line with observed data.

Here, we propose Mutual Hazard Networks (MHN). Rather than grouping events into pathways, MHN model both co-occurence and mutual exclusivity by direct interactions between events. MHN characterize events by a combined rate of occurance and spontaneous fixation and by multiplicative effects they exert on the rates of successive events. These effects can be cyclic and greater or less than one, i.e., promoting or inhibiting. We provide formulas for the log-likelihood of MHN and its gradient, and an implementation that is computationally tractable for systems with up to 25 events on a standard workstation and for larger systems on an HPC infrastructure.

## 2 Methods

### 2.1 Mutual Hazard Networks

We model tumour progression as a continuous time Markov process {*X*(*t*), *t* ≥ 0} on all 2^*n*^ combinations of a predefined set of *n* events. Its state space is *S* = {0, 1}*^n^*, where *X*(*t*)_*i*_ = 1 means that event *i* has occurred in the tumour by age *t*, while *X*(*t*)_*i*_ = 0 means that it has not.

We assume that every progression trajectory starts at a normal genome *X* (0) = (0, …, 0)^*T*^, accumulates irreversible events one at a time, and ends at a fully aberrant genome *X*(*∞*) = (1, …, 1)^*T*^. Observed tumour genomes correspond to states at unknown intermediate ages 0 < *t* < ∞ and typically hold both 0 and 1 entries.

Let *Q* ∈ ℝ^2^*n*^×2^*n*^^ be the transition rate matrix of this process with respect to a basis of *S* in lexicographic order (Fig. 1, left). An entry

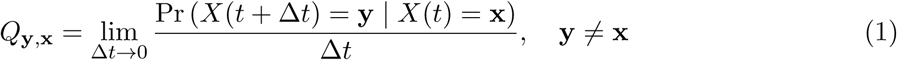

is the rate from state x ∈ *S* to state y ∈ *S*, and diagonal elements are defined as *Q*_x, x_ = −Σ_y≠x_ *Q*_y, x_ so that columns sum to zero. *Q* is lower triangular and has non-zero entries only for transitions between pairs of states x = (…, *x*_*i*−1_, 0, *x*_*i*+1_, …)^*T*^ and y = x_+*i*_:= (…, *x*_*i*−1_, 1, *x*_*i*+1_, …)^*T*^ that differ in a single entry *i*.

**Figure 1:**
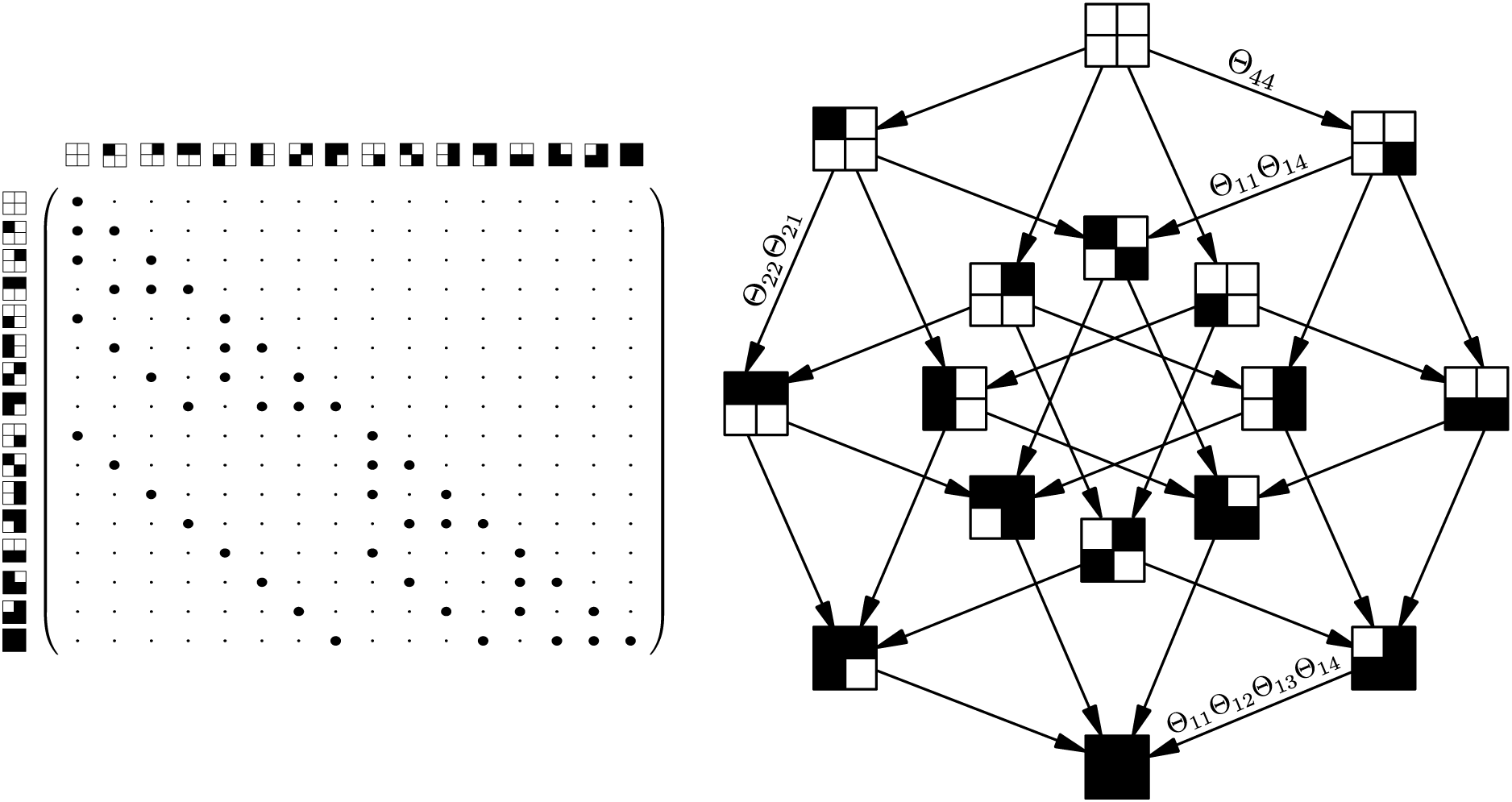
(Left) Transition rate matrix *Q* for the Markov process *X* with *n* = 4. It is lower triangular because events are irreversible, and sparse because events accumulate one at a time. (Right) Parameterization *Q*_e_ of the Markov process by a Mutual Hazard Network.

Our aim is to learn for each event *i* how its rate of fixation *Q*_x+*i*, x_ depends on preceding events in x. It is, however, impractical to treat all entries in *Q* as free parameters because of its exponential size. Instead we parameterize *Q* by a *Mutual Hazard Network* which is a smaller *n* × *n* matrix Θ with positive real entries. It restricts rates in *Q* to the functional form

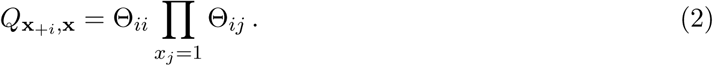

Here, Θ_*ii*_ is the baseline rate of spontaneous fixation of event *i* when it occurs before any other event. Θ_*ij*_ is the multiplicative effect by which a preceding event *j* in x modulates the rate of *i* (Fig. 1, right).

### 2.2 Parameter Estimation

A dataset 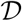 of tumours defines an empirical probability distribution on *S*. It can be represented by a vector 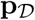 of size 2^*n*^, where an entry 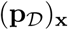 is the relative frequency of observed tumours with state x in 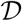.

At *t* = 0 tumours are free of any events, so the Markov process *X* starts with the initial distribution p_*∅*_:= (100%, 0%,*…,* 0%)^*T*^, which then evolves according to the parameterized rate matrix *Q*_Θ_. If all tumours had been observed at a common age *t*, 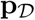 could be modelled as a sample from the transient distribution

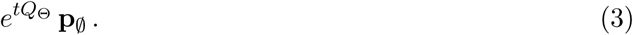

Since the tumour age is usually unknown, we follow Gerstung et al. (2009) and consider *t* to be an exponential random variable with mean 1. Marginalizing over *t* yields

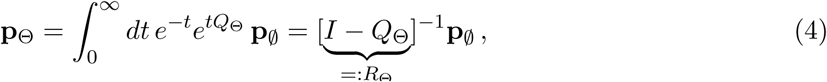

and the marginal log-likelihood score of Θ given 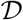 is

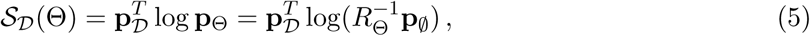

where the logarithm of a vector is taken component-wise.

When optimizing 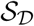 with respect to Θ we are especially interested in networks that can be easily visualized and interpreted, i.e., where many events do not interact and off-diagonal entries Θ_*ij*_ are exactly 1. To this end, we penalize the score with a sparsity-promoting regularization term,

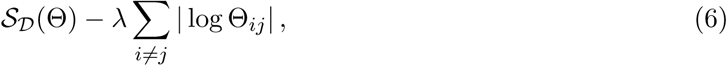

where λ is a tuning parameter. We will optimize this expression using the Orthant-Wise Limited-Memory Quasi-Newton algorithm (Andrew and Gao, 2007). This general-purpose optimizer takes care of the non-differentiability introduced by the regularization term, while only requiring a closed form for the derivatives 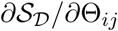 with respect to each parameter.

From the chain rule of matrix calculus we have

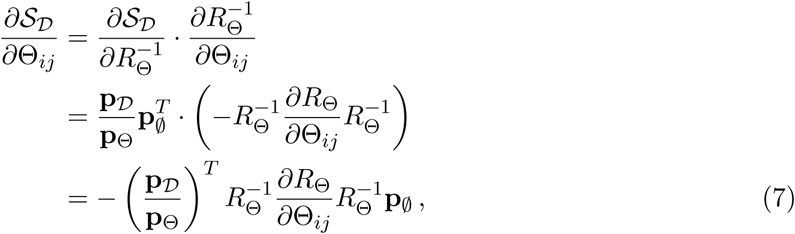

where *·* is the Frobenius product and the ratio 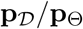 is computed component-wise.

### 2.3 Efficient implementation

To compute the score in equation (5) and its gradient in equation (7) we must solve the exponentially sized linear systems [*I* − *Q*_Θ_]^−1^ p_∅_ and 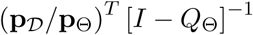. To this end, we employ the (left) Kronecker product which is defined for matrices *A* ∈ ℝ^*k×l*^ and *B* ∈ ℝ^*p×q*^ as the block matrix

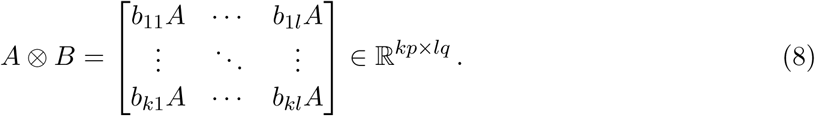

We follow the literature on structured analysis of large Markov chains (Buchholz, 1999; Amoia et al., 1981) and write the transition rate matrix *Q*_Θ_ as a sum of *n* such Kronecker products,

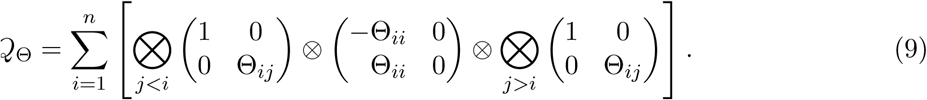

Here, the *i*-th term in the sum is a sparse 2*^n^* × 2^*n*^ matrix consisting of all transitions that introduce event *i* to the genome. It corresponds to a single subdiagonal of *Q*_Θ_, together with a negative copy on the diagonal to ensure that columns sum to zero (Fig. 2). The benefit of this compact representation is that matrix-vector products can be computed in 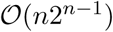 rather than 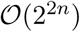 without holding the matrix explicitly in memory (Buis and Dyksen, 1996). We split *R*_Θ_ = *I* − *Q*_Θ_ into a diagonal and strictly lower triangular part,

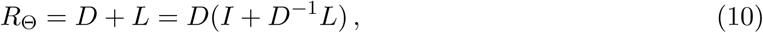

and use the nilpotency of *D*^−1^*L* to compute

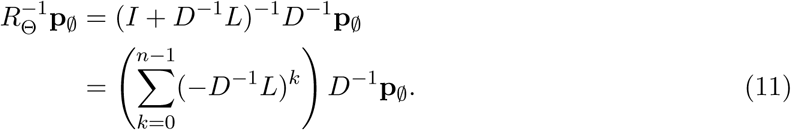

**Figure 2:**
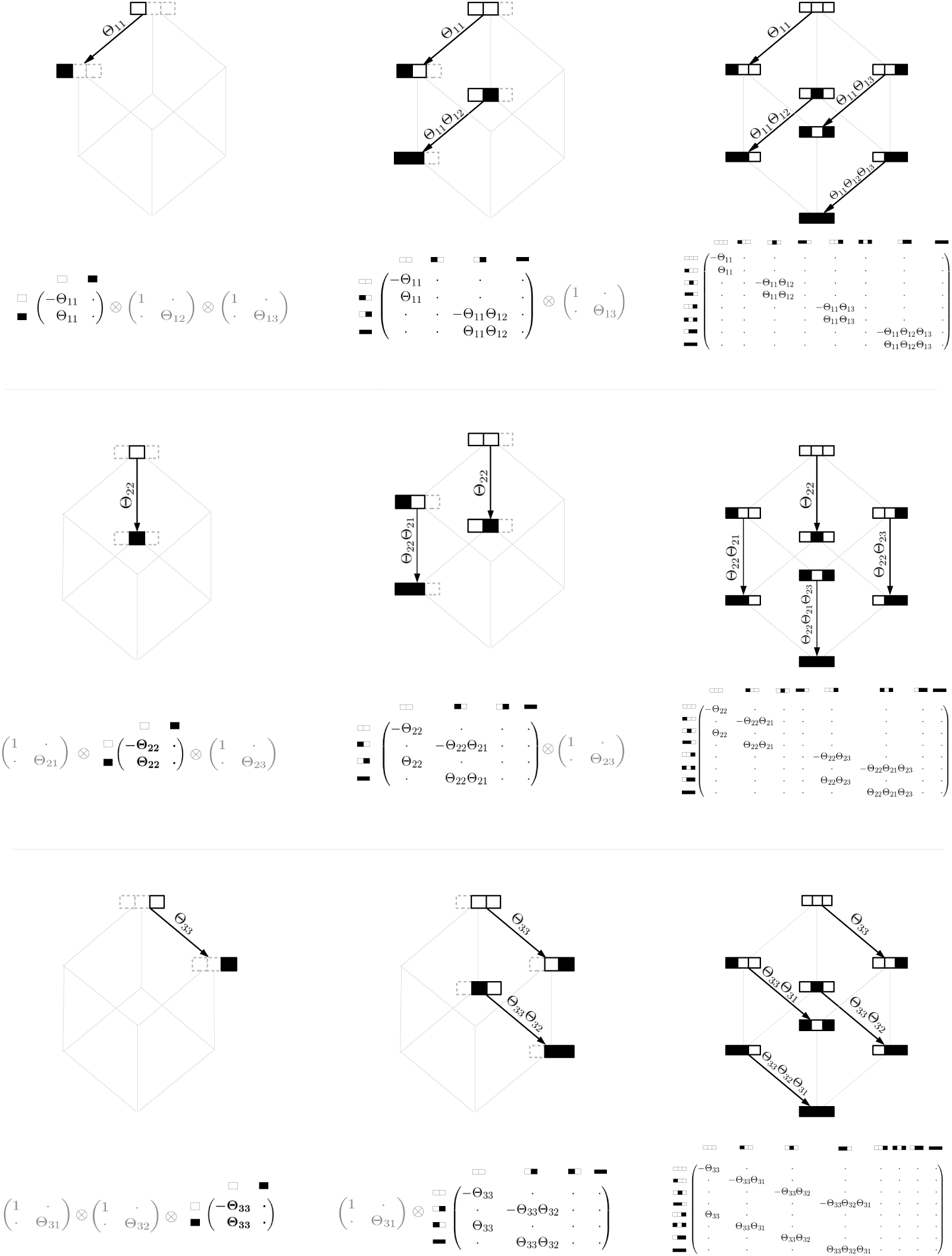
Illustration of *Q*_Θ_ represented as a sum of Kronecker products for *n* = 3 in equation (9). The *i*-th row corresponds to the *i*-th term in the sum and contains all transitions that introduce event *i* to the genome. A row is read from left to right and shows how the Kronecker product successively describes all possible transition rates that can arise due to multiplicative interactions with other events. The first highlighted Kronecker factor describes the two possible states of event *i* and a transition with base rate Θ_*ii*_. Each subsequent Kronecker factor that is multiplied from the left or from the right appends the two states of the corresponding event *j* to all previously modelled states. This doubles the number of modelled states, where one half lacks the event *j* and retains their previous transition rates, while the other half has *j* present, which modulates their transition rates by the factor Θ_*ij*_.

## 3 Results

### 3.1 Simulations

We tested in simulation experiments how well an MHN of a given size can learn a probability distribution on *S* when trained on a given amount of data. We ran 100 simulations for each of several sample sizes 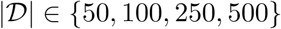 and number of events *n* ∈ {10, 15}.

In each simulation run, we chose a ground truth model Θ with *n* possible events. A random half of its off-diagonal entries were set to 1 (no interaction) and the remaining entries were drawn from a standard log-normal distribution. We then generated a dataset of size 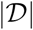 from this model and trained on it another model 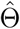 by optimizing expression (6). We chose a common regularization parameter for all 100 simulation runs, which we found to be roughly 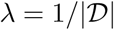 through validation on separate datasets of each sample size. We then assessed the reconstructed model 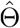 by the Kullback-Leibler (KL) divergence from its probability distribution to the distribution of the true model Θ,

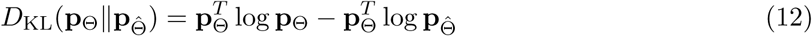

The median KL divergence, as well as its variance over the 100 simulation runs, improved with larger training datasets and reached almost zero (Fig. 3).

**Figure 3:**
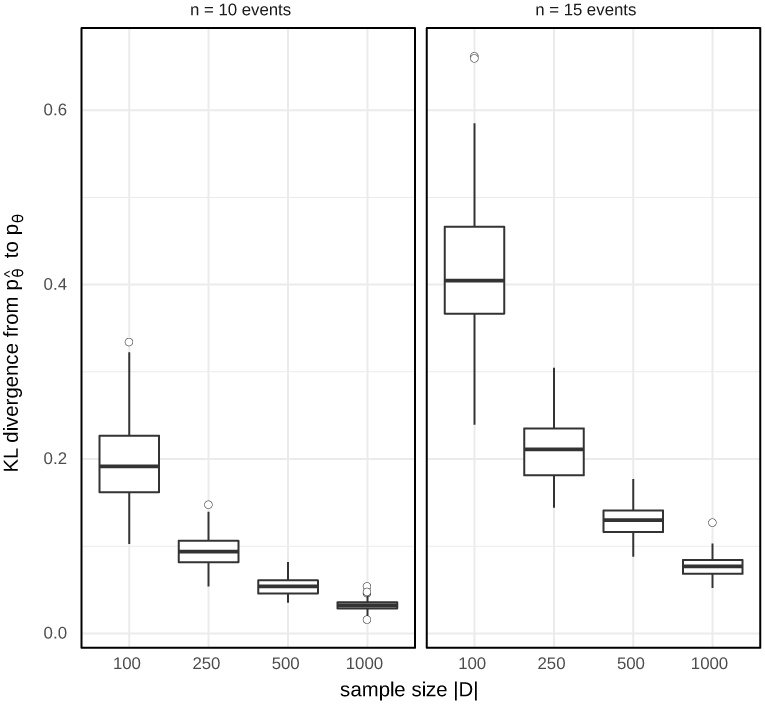
Model fit for different sample sizes in simulation experiments.

Next, we tested the performance of our implementation. MHN was written in R, and its performance-critical parts were implemented in C (using the R package inline) to avoid unnecessary memory-copy operations. We made explicit calls to BLAS routines and compiled R to use the Intel MKL library for vectorized and threaded matrix and vector operations. Fig. 4 shows the runtime of a single gradient step for random and dense Θ on a Dell OptiPlex 9020 workstation with 8GB RAM and an Intel^®^ Core^TM^ i5-4590 CPU. The runtime was about 1 minute for *n* = 20 and scaled exponentially with *n* as expected.

**Figure 4:**
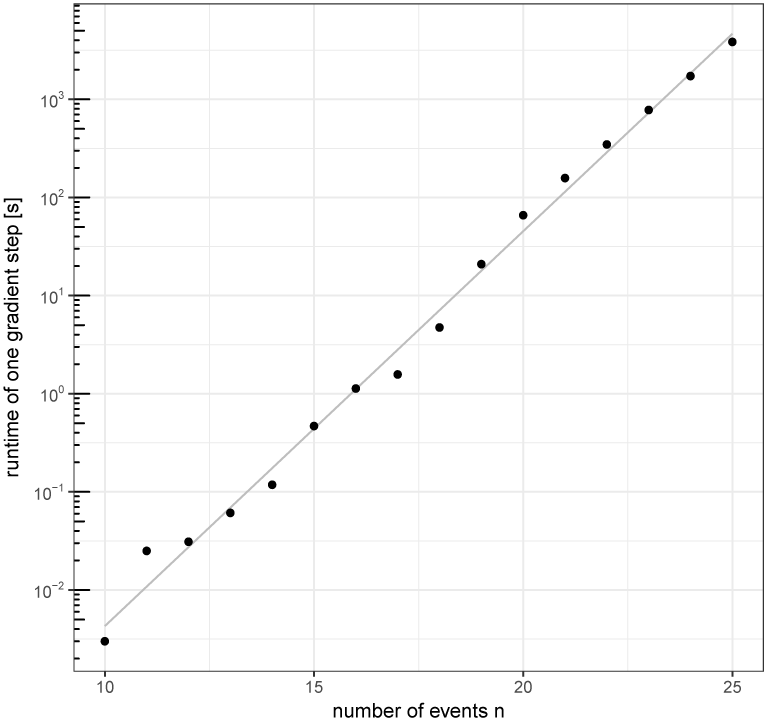
Runtime of a single gradient step for random and dense Θ.

### 3.2 Application to Cancer Progression Data

#### 3.2.1 Comparison to Conjunctive Bayesian Networks

We tested our method and first compared it to Conjunctive Bayesian Networks (CBN) on three cancer datasets that were previously used by Gerstung et al. (2009). They were obtained from the Progenetix molecular-cytogenetic database (Baudis and Cleary, 2001) and consist of 817 breast cancers, 570 colorectal cancers, and 251 renal cell carcinomas. The cancers are characterized by 10, 11, and 12 recurrent copy number alterations, respectively, which were detected by comparative genomic hybridization (CGH).

We trained MHN on all three datasets (see supplementary material) and compared them to the CBN given in Gerstung et al. (2009), which provide log-likelihood scores in-sample. Since the in-sample scores of MHN are biased by their greater flexibility, we also provide their average log-likelihood scores in 5-fold cross-validation. To avoid a nested loop for tuning the sparsity parameter λ we set it to a fixed value of 0:01. Despite these handicaps, MHN compared favourably on all three datasets (Table 1)

**Table 1:** Log-likelihood scores

| dataset | (cross-validated) | (in-sample) |  |
| --- | --- | --- | --- |
|  | MHN | CBN | MHN |
| Breast cancer | -5.68 | -5.73 | -5.62 |
| Colorectal cancer | -5.66 | -5.79 | -5.62 |
| Renal cell carcinoma | -5.04 | -5.13 | -4.87 |

#### 3.2.2 Comparison to pathTiMEx

Next, we compared MHN to pathTiMEx on a glioblastoma dataset from The Cancer Genome Atlas (Cerami et al., 2012) which was previously used in Cristea et al. (2017), see Fig. 5. The data consist of 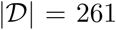 tumours characterized by 486 point mutations (M), amplifications (A), or deletions (D). We focus on *n* = 20 of these events which were pre-selected by pathTiMEx using the TiMEx algorithm (Constantinescu et al., 2015).

**Figure 5:**
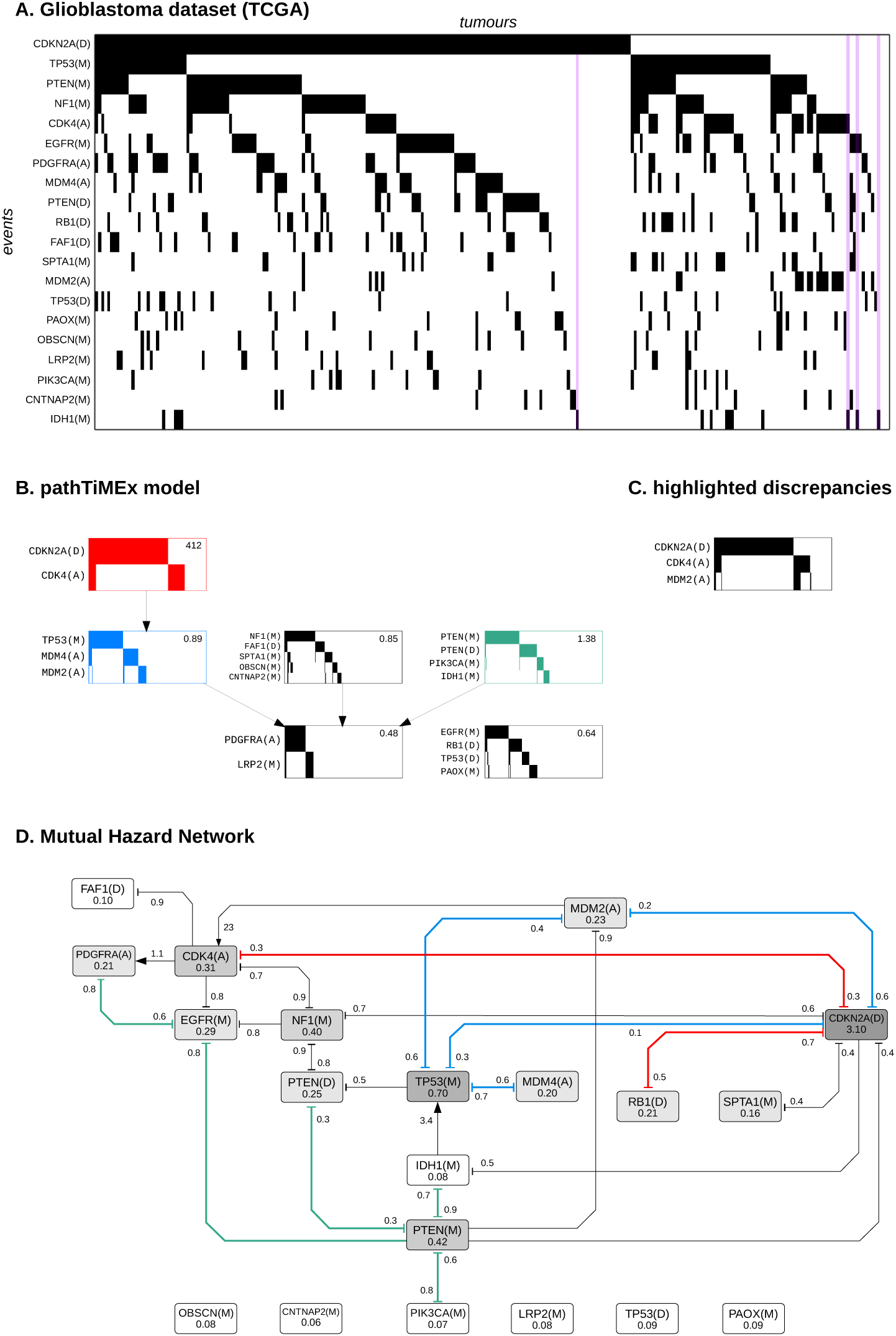
(A) Glioblastoma dataset from TCGA, where rows show events sorted by frequency and columns show tumours sorted lexicographically. The purple stripes highlight tumours which have IDH1(M) but lack TP53(M). (B) PathTiMEx model inferred in Cristea et al. (2017). It simultaneously divides the dataset into pathways, i.e., into mutually exclusive groups of events and learns a CBN of these pathways. The CBN considers a pathway altered if at least one of its constituent events has occured. A pathway alteration fixates at the rate given in the upper right-hand corner once all its parent pathways in the CBN have been altered. (C) Highlighted discrepancies between the data and the pathTiMEx model due to its assumption of interchangeable events. Although *CDKN2A(D*) and *CDK4(A*) were grouped into the same pathway, *CDKN2A(D*) is negatively associated with *MDM2(A*) in the data while *CDK4(A*) is positively associated with it. (D) Mutual Hazard Network, where nodes show the base rates Θ_*ii*_ and edges show the multiplicative interactions Θ_*ij*_. Similarities to pathTiMEx are highlighted in colour and roughly correspond to the signaling pathways Rb, p53, and PI(3)K (red, blue, and green).

We trained MHN as above for 100 iterations, which achieved a log-likelihood score of -7.70 in-sample and a score of -7.97 in 5-fold cross-validation. While pathTiMEx does not yield a directly comparable log-likelihood score, it quantifies discrepancies between model and data by considering the data to be corrupted by noise, each event in a tumour being independently flipped with probability ɛ. PathTiMEx estimated this noise parameter as 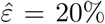, from which we gauge an upper bound on its log-likelihood score as follows: even a hypothetical model that learns the data distribution 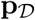 perfectly but assumes a level of noise

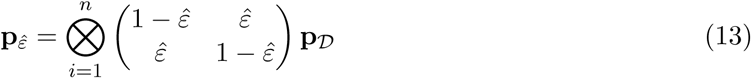

achieves only a score of 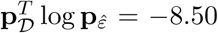 in-sample, which is less than the cross-validated score of MHN.

Nevertheless, MHN largely agreed with pathTiMEx on the three most mutually exclusive groups of events. They broadly correspond to the signaling pathways Rb, p53, and PI(3)K (red, blue, and green in Fig. 5) which regulate cell cycle progression, apoptosis, and proliferation and are well known to be compromised in glioblastoma (McLendon et al., 2008). Where the models differ, MHN more closely matches the literature and additionally included *RB1(D*) in the Rb pathway, *EGFR(M*) and *PDGFRA(A*) in the PI(3)K pathway, and *CDKN2A(D*) in the p53 pathway. It also correctly identified the fact that the Rb and p53 pathways overlap and that both involve *CDKN2A(D*) which codes for two different proteins p16^INK4a^ and p14^ARF^ in alternate reading frames.

Notably, MHN inferred that the rare event *IDH1(M*) promotes the more common event *TP53(M*). This is further illustrated in Fig. 6 which shows the most likely chronological order of events for all 261 tumours. Each of their 193 distinct states is represented by a path that starts at the root node and terminates at either a leaf node or an internal node with a black outline. As can be seen in the lower left, all tumours that contain *IDH1(M*) are located on a common branch and thus share an early mutation history initiated by *IDH1(M*). This interpretation is in line with the fact that *IDH1(M*) is considered a defining attribute of the Proneural subtype of gliobastoma which is clinically distinct and also associated with *TP53(M*) (Verhaak et al., 2010). It is further supported by independent data from consecutive biopsies of gliomas where *IDH1(M*) in fact preceded *TP53(M*) (Watanabe et al., 2009).

**Figure 6:**
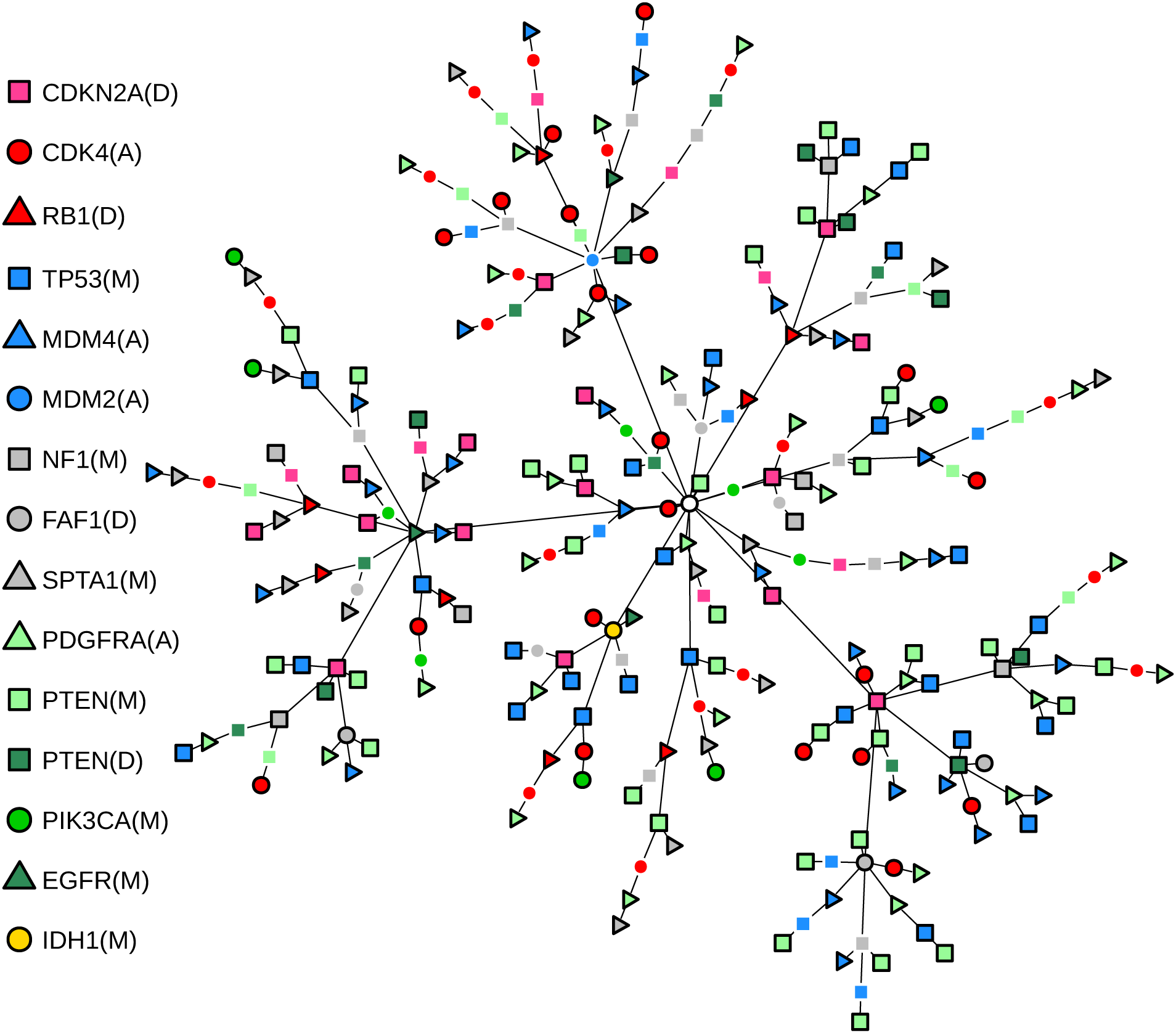
Maximum likelihood paths through the state space *S* from the starting state to each observed tumour state. They were computed from the time-discretized transition rate matrix 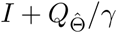, where γ is the greatest absolute diagonal entry of 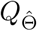.

## 4 Discussion

We presented Mutual Hazard Networks, a new framework for modelling tumour progression from cross-sectional observations. MHN build on previous work (Beerenwinkel et al., 2007; Hjelm et al., 2006; Raphael and Vandin, 2015; Cristea et al., 2017) and extend it in multiple ways: (a) MHN naturally account for any form of epistatic interactions including inhibition, promotion, and cycles. (b) MHN do not rely on a hard grouping of events into pathways, hence allowing for overlap or cross-talk. (c) MHN do not rely on the interchangeability assumption for mutually exclusive events. In other words: they do not assume that the future progression of a tumour is independent of which particular gene in a pathway was actually affected by a mutation.

These issues matter, as has become clear in the application to glioblastomas: (a) MHN detected several inhibiting edges as well as cyclic interactions that remained obscure in acyclic models, (b) MHN naturally resolved the role of *CDKN2A*, which is involved in at least two pathways (McLendon et al., 2008), and (c) MHN uncovered that the interchangeability assumption does not hold for the *CDK4(A)-CDKN2A(D*) group (Fig. 5C).

Our proposed implementation of the MHN learning algorithm has a space and time complexity that is exponential in the number of events *n*. In practice, we saw limits at *n* = 25 on a standard workstation. Modern cancer datasets report hundreds of recurrent mutations, and the question arises whether MHN can deal with them. In fact we believe that MHN is competitive with other algorithms also for these large datasets, because interactions between low-frequency events cannot be resolved reliably at all. For example, in the glioblastoma dataset, the rare events *OBSCN(M), CNTNAP2(M), LRP2(M), TP53(D),* and *PAOX(M*) remained unconnected to the rest of the network. In other words, the evidence for possible interactions was so low that it could not compensate for the L1-costs of an additional edge. These are limitations in the data itself and not in computation times.

An interesting novelty of MHN are the spontaneous occurrence/fixation rates _*ii*_. The event pair *IDH1(M*) and *TP53(M*) was instructive for understanding their role.*IDH1* mutations were infrequent in the glioblastomas compared to *TP53* mutations. Moreover, 10 out of 14 *IDH1(M*) positive glioblastoma also showed a *TP53* mutation. We see at least two alternative explanations for this noisy subset pattern: (1)*TP53* mutations are needed for *IDH1* mutations to occur. (2) *TP53(M*) has a much higher spontaneous rate than *IDH1(M*) explaining that it is more frequent, and moreover, an *IDH1* mutation strongly increases the rate of a *TP53* mutation, explaining why so many *IDH1(M*) positive glioblastoma were also positive for *TP53(M*). While both scenarios explain the noisy subset pattern, they disagree with respect to the chronological order of events. In (1) the *TP53* mutation precedes the *IDH1* mutation, while in (2) the events occur in reverse order. MHN decided for explanation (2) and is endorsed by independent data from consecutive biopsies (Watanabe et al., 2009). Where in the training data was the evidence in favour of (2)? We found it in the four *IDH1(M*) positive /*TP53(M*) negative cases (Fig. 5A, purple). All of them had at most one mutation in addition to *IDH1(M*), which is in line with (2) but not with (1).

In summary, we introduced a new, very flexible framework for tumour progression modelling that naturally accounts for cyclic interactions between events.

## Acknowledgements

This work was funded by DFG grants FOR 2127 and SFB/TRR-55. We thank Daniel Richtmann and Stefan Hansch for helpful discussions.

